# Stress induced Differential Expression of THAP9 & THAP9-AS1 in the S-phase of cell cycle

**DOI:** 10.1101/2021.02.11.430738

**Authors:** Vasudha Sharma, Prachi Thakore, Meena Krishnan, Sharmistha Majumdar

## Abstract

Transposable elements function as one of the major effectors in response to biological or environmental stress. Under normal conditions, host organisms deploy epigenetic and post-transcriptional machinery (histone modifications, chromatin remodelers, long non-coding RNAs (lncRNAs)) at the TE sites to contain their mobility. But many a times, the chromatin architecture undergoes TE induced changes under the effect of stress that in turn might lead to unprecedented gene expression. LncRNAs are emerging as a crucial tool in the regulation of TEs. TEs possess remarkable abilities to respond in the face of stress, ranging from undetected mutations to changing the regulatory landscape of the host. Although the relationship between stress response and TE activation/deactivation is well acknowledged but our understanding of the mechanism of regulation remains poor.

This study focuses on the gene expression of THAP9, a domesticated transposon and lncRNA THAP9-AS1 (THAP9-anti sense1), which form a sense and anti-sense gene pair with a promoter overlap of approximately 350bp. The two genes exhibit different patterns of gene expression under different types of stresses in the S-phase of the cell cycle. THAP9-AS1 is always upregulated under stress whereas THAP9 exhibits both downregulation and upregulation in different stresses. Both THAP9 and THAP9-AS1 exhibit a periodic gene expression throughout the S-phase which is a characteristic of cell cycle regulated genes.

## Introduction

Transposons or transposable elements (TEs) are highly dynamic sequences that can move from one genomic region to another (mobile DNA) without the need of homology between donor and target sequences. Owing to their self-propagatory nature, these sequences were labelled as ‘parasitic’ and ‘selfish’ in nature. A significant portion (3-50%) of the genomes of most species is contributed by TEs (1, 2). Decades of research has led to a paradigm shift in biology where it is now a well accepted fact that transposons are not parasitic but essential players in creating genomic diversity leading to evolution (3, 4). Many TEs have been reported to be domesticated and repurposed by the host organisms for their own benefit, eg, CENPB, Rag1, Rag2, SETMAR, THAP9 (5).

TEs were first described in 1984 by McClintock in her seminal study, reporting that the expression of TEs change in response to genomic insults and the outcomes of transpositions can be detrimental to the structure, function and evolution of the genome (6). To maintain cellular homeostasis, the transposon sequences are controlled from jumping around the genome by various epigenetic mechanisms (7) . Heavy repressive modifications of Histone (H3K9 methylation) and DNA methylation are often observed in TE rich regions in several organisms and mutations resulting in loss of methylation causes upregulation of TEs (8, 9). ATP-dependent chromatin remodelers (SWI/SNF) use their ATPase domain to hydrolyze ATP to modify nucleosomes to silence TEs (10). KRAB-ZFPs (Kruppel-associated box zinc-finger proteins), the largest class of mammalian DNA binding transcription factors, silence TEs by recruiting co-repressor proteins (11). piRNAs (PIWI interacting RNAs) carry inactive remnants of TEs and are known to repress TE activity in the germline (12).

Many lncRNAs or long non coding RNAs (length >200 nucleotides) are located proximal to certain TEs and may contribute to their regulation (13, 14). lncRNA-mediated TE repression may have evolved as a mechanism to counteract transposon-induced genomic instability (e.g., oncogenic translocations in several tumors) (15). Interestingly, about two-thirds of total human lncRNAs contain TE sequences that are inactive or remnants of active TEs (13).For example, many lncRNAs have LINE, Alu, ERVs and SINE sequences embedded in them and mutations in these regions may lead to lethal diseases (16).

Several studies have reported the activation or repression of TEs in response to stress(1). When stimulated by external stress, the cellular homeostasis gets disrupted, often leading to epigenetic changes which in turn might cause reactivation of transposons resulting in insertional mutagenesis (17). For example, under stress conditions, LINE-1 elements get activated upon release of SIRT-6 protein binding. Stress has also been reported to silence or deactivate transposons. For example, in yeast, Ty3 transposons get repressed under heat stress (18). Stress can also increase histone methylation in a tissue specific manner resulting in the silencing of various families of activated transposons (19). The number of Class II (DNA) transposons reported to be regulated under stress conditions is far less than Class I transposons (7, 20).

The activation of TEs under stress can be both beneficial or deleterious for the host (7). This is because the actual relationship between stress and TE expression is quite complex and varies depending upon the type of stress, type of organism and the epigenetic/genetic/metabolic changes drawn out by the stress (7). The regulation may be temporal or spatial wherein certain TEs can be turned on at specific stages of embryonic development or in specific cell types or tissues (7). Since stress induces both the defence response and the activation/deactivation of certain TEs within a host, it is hard to decipher the cause-effect relationship between the TEs and stress. Stress often results in double stranded DNA breaks which are detected and repaired at the DNA-integrity checkpoints of the cell cycle, by NHEJ (Non homologous end joining) when they occur in early S/G_1_ phase or Homologous repair when they occur in late S/G_2_ phase (21, 22). Interestingly, p53, which is known for its role in maintaining genomic stability during external stress and is mutated in almost all cases of cancer (23–25), may directly repress certain transposons (26).

We are interested in THAP9, a recently discovered human DNA transposase, which is homologous to the widely studied Drosophila P-element transposase (27). The THAP9 protein shares 40% similarity to the P-element transposase, and probably lacks the ability to transpose due to the absence of terminal inverted repeats and target site duplications. THAP9 is present as a single copy in the human genome and its function has not been discovered yet. Despite losing the hallmarks of a transposon, it has retained its catalytic activity (27, 28). Moreover it is not known if and how THAP9 is regulated at the cellular level.

The human THAP9 gene encodes for 6 transcripts, out of which only one encodes for the full-length transposase protein. THAP9-AS1 (THAP9 antisense) is a newly annotated lncRNA coding gene by Ensembl that encodes for 12 long non-coding RNAs (29, 30). The expression of THAP9 and THAP9-AS1 is controlled by overlapping promoters on opposite strands, as curated by FANTOM database (31, 32) (Fig 1). Antisense transcripts and their corresponding sense transcripts often show inverse expression; thus suggesting that one regulates the other (33, 34). There have been recent reports where the THAP9-AS1 lncRNA has been implicated in pancreatic cancer, septic shock and neutrophil apoptosis (35–37). This has piqued our interests in studying the relationship between THAP9 and THAP9-AS1.

**Fig 1:**
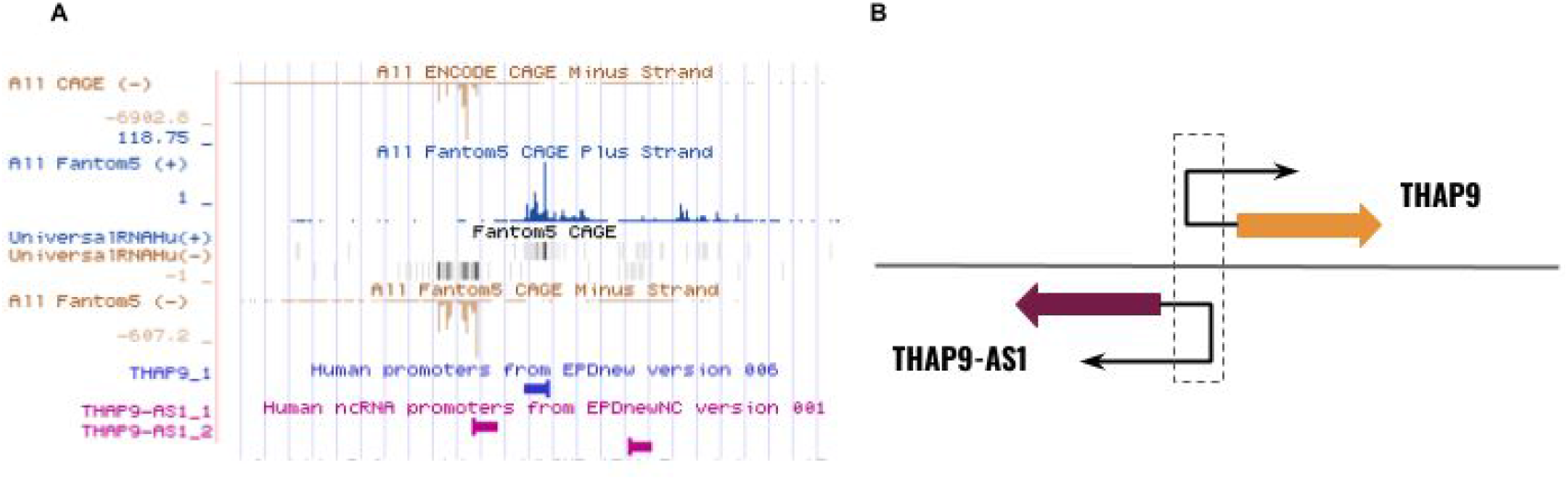
(A) THAP9 and THAP9-AS1 are controlled by different promoters on (+) and (−) strands. Promoter upstream of THAP9 (on (+) strand (shown in blue)) and THAP9-AS1 gene (on (−) strand (shown in pink)) as reported by CAGE data from FANTOM database (31). (B) Representative figure of overlapping promoters of THAP9 and THAP9-AS1.

In this study, we have investigated the effect of stress on THAP9 expression. We have subjected a synchronized population of HEK293T cells in S-phase to various stress conditions, namely, heat shock, genotoxic, osmotic and oxidative stress and quantified THAP9 and THAP9-AS1 RNA expression by qPCR. This study reports a detailed quantitative analysis of THAP9 and THAP9-AS1 gene expression in the S-phase under various stress conditions.

## Materials and methods

### Reagents

HEK293T cell lines, DMEM (HyClone) supplemented with 10% Fetal Bovine Serum (Gibco), TRIzol™ Reagent (Invitrogen), High-Capacity cDNA Reverse Transcription Kit (Applied Biosystems™ 4368814), mouse anti-cyclin D1 monoclonal antibody (Invitrogen AHF0082), mouse anti-cyclin E monoclonal antibody (Invitrogen MA514336), mouse anti-cyclin A2 monoclonal antibody (Invitrogen MA1180), mouse anti-actin monoclonal antibody (Invitrogen MA1744), KAPA^R^ SYBR Fast Universal kit (Sigma KK4602)

### Cell synchronization by double thymidine block

0.3 × 10^5^ HEK293T cells (grown in DMEM supplemented with 10% fetal bovine serum under standard tissue culture conditions) were plated per well of 6-well plates and incubated for 12 hours at 37*C and 5% CO_2_. To get a homogenous population of cells at G_1_/S phase boundary, thymidine (Sigma T9250) was added to a final concentration of 2mM and incubated for 18 hours at 37*C. Cells were then washed with 1 ml pre-warmed 1X PBS and incubated with fresh media for 9 hours at 37*C. This was followed by second dose of thymidine at a final concentration of 2 mM and incubation for 18 hours at 37*C.

### Assessment of cell synchrony

The cell lysates collected at 0h, 2h, 4h, 6h, 8h in the S-phase were run on 10% denaturing SDS PAGE in a running buffer (1X Tris- Glycine SDS). PageRuler™ Prestained Protein Ladder (10 to 180 kDa, ThermoFisher Scientific; 26616) was used as a molecular size marker and the samples were transferred to PVDF membrane by electroblotting using a standard gel transfer system. The membrane was blocked to remove nonspecific binding using 3% (w/v) skimmed milk in Tris-Buffered Saline with Tween-20 (TBST). Membranes were washed with 1X TBST and incubated overnight at 4°C with 1:1000 dilution of primary cyclin-D, 2ug/ml cyclin-E, 2ug/ml cyclin-A and 1:1000 beta-actin primary antibody followed by incubation with 1:5000 dilution of HRP- coupled secondary antibodies for 2h at room temperature and detected using enhanced chemiluminescence (Pierce, PI32106) on a Bio- Rad Gel Documentation system.

### Stress Treatments

#### Heat stress

After the second thymidine block,the cells were subjected to heat at 42-43 degrees for 20 min to induce heat shock. After 20 minutes, the media was changed and cells were collected at 0 hour, 2 hour, 4 hour, 6 hour, 8 hour to cover the entire span of S-phase.

#### Genotoxic stress

Cells were kept under UV light for 20 min after double thymidine block. This was followed by addition of fresh media and collection of cells at 0h, 2h, 4h, 6h, 8h.

#### Osmotic stress

Different concentrations of DMEM were prepared by using dissolving powdered form of the medium (Himedia AT186) to prepare 10X and 0.1X solutions supplemented with 10% fetal bovine serum and 1% antibiotic (Penicillin-Streptomycin-Glutamine Gibco 10378-016). After the double thymidine block, cells were grown in different osmotic conditions for 30 min followed by replacement with fresh media and collection of cells at 0h, 2h, 4h, 6h, 8h.

#### Oxidative stress

Hydrogen peroxide (Merck 18304) was used in varying concentrations (final concentrations of 10uM, 25uM and 50uM) to induce oxidative stress in HEK293T cells. Following the double thymidine block, cells were incubated in media containing hydrogen peroxide for 30 min. After incubation fresh media was added to the cells and cells were collected at 0h, 2h, 4h, 6h and 8h.

### RNA Isolation, cDNA preparation

After the respective stress treatments, cells were isolated at their specific time points (0h, 2h, 4h, 6h, 8h) by scraping each well in 1ml Trizol after a wash with 1X PBS. The cell lysate was mixed thoroughly and transferred to the eppendorf tube and incubated for 5 minutes. 250ul of chloroform (Sigma 2566) was added and vortexed for 15-20 secs and kept at room temperature for 5 min, It was centrifuged at 10,000rpm for 5 min. The top aqueous layer was transferred carefully to a fresh tube and 550ul of isopropanol (Fisher scientific 26897) was added and mixed by gently inverting the tube 5-6 times and kept at room temperature for 5 min. The tubes were centrifuged at 14,000rpm for 30 min. Supernatant was decanted and 1ml of 75% ethanol prepared in DEPC (Sigma D5758) treated water was added, mixed gently and centrifuged at 9,500rpm for 5 min. The pellet was air- dried and resuspended in 15-20ul of DEPC water. The RNA was quantified and the samples with optical density 260/280 greater than to 1.8 were used for cDNA preparation.

cDNA synthesis was carried out as per manufacturer’s protocol using High-Capacity cDNA Reverse Transcription Kit (Applied Biosystems™ 4368814). The synthesized cDNA was quantified and diluted to a final working concentration of 50ng for qPCR.

### qPCR

qPCR reaction was set up as per manufacturer’s instructions (Applied Biosystems™ 7500) with 50ng cDNA. The primers (F= forward, R=reverse) used for amplification are as follows: THAP9 (F:5’-GGGGTTTATGTGCTTTGGTCTTGGAAAAC-3’, R:5’-GCAATCTGTTGACTAGAAG-3’) THAP9-AS1 (F:5’-CACAATCTGGCGCCATCG-3’, R:5’-CCTTATTTCCTTCATTGTGGCAAAG-3’) P53 (F:5’-CCTCAGCATCTTATCCGAGTGG-3’, R:5’-TGGATGGTGGTACAGTCAGAGC-3’) HSP70 (F:5’-ACCTTCGACGTGTCCATCCTGA-3’, R:5’-TCCTCCACGAAGTGGTTCACCA-3’) SOD1 (F:5’-GTAGTCGCGGAGACGGGGTG-3’, R:5’-GAGGCCTGGCGGGCGAC-3’) HPRT1 (F:5’-CATTATGCTGAGGATTTGGAAAGG-3’, R:5’-CTTGAGCACACAGAGGGCTACA-3’)

### Data Analysis

<u>Fold change of each gene</u> was measured by comparative C_T_ method (also known as 2^−ΔΔCt^ method) using following equation (38):

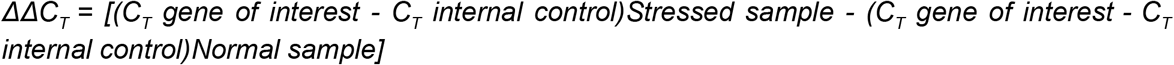

<u>Gene expression under normal conditions</u> was calculated as 2^−ΔCt^ where ΔC_T_ is the difference between C_T_ values of gene of interest and HPRT.

Two-tailed, unpaired Student’s t test was used to evaluate expression differences between gene of interest and internal control gene (HPRT1). P values of <0.05 were considered statistically significant

## Results

### Expression of THAP9 and THAP9-AS1 increases throughout the S-phase in response to genotoxic stress

External or internal stress can stimulate changes in the activity of transposable elements(1). No information is available for the stress response of THAP9 and THAP9-AS1. Approximately 1100 lncRNAs are known to be enriched in S phase (39). Although the exact role of this upregulation of lncRNAs in S phase remains undiscovered but the most prevalent hypothesis is that they express in response to DNA damage during the cell cycle (40).

Instead of recording stress response at random time points, we followed the recent studies of lncRNAs expression in the S-phase for studying the stress induced change in gene expression of THAP9 and THAP9-AS1 (39, 40). For quantifying the expression of THAP9 and THAP9-AS1, we first synchronized the cells at the G_1_/S phase border of the cell cycle followed by UV irradiation to induce genotoxic stress response in the S-phase which typically lasts for 6-8 hours in mammalian cells. We then measured THAP9 and THAP9-AS1 transcripts in early S-phase (0-4h), mid S-phase (4-6h) and late S-phase (6-8h). To ensure that the cells are in S-phase we probed them for the expression of S-phase specific cyclins (cyclin A and cyclin E) (Fig 2). To further make sure that the stress treatment was effective, we quantified the cells for P53 transcripts which is a positive biomarker for genotoxic stress. The expression of P53 shows an oscillatory nature throughout the S-phase which indicates that the stress treatment has worked (41). THAP9-AS1 was upregulated under UV stress and followed a similar pattern to that of P53, wherein the expression of both the genes falls at 2h and then increases consistently until the end of the S-phase (p<0.01) (Fig). The highest expression of THAP9-AS1 is recorded at 8 hour which is approximately 8 fold higher than the recording at 0 hour. qPCR results also revealed a consistent upregulation in the expression of THAP9 transcripts, highest during the late S-phase (p<0.01) (Fig 3).

**Fig 2:**
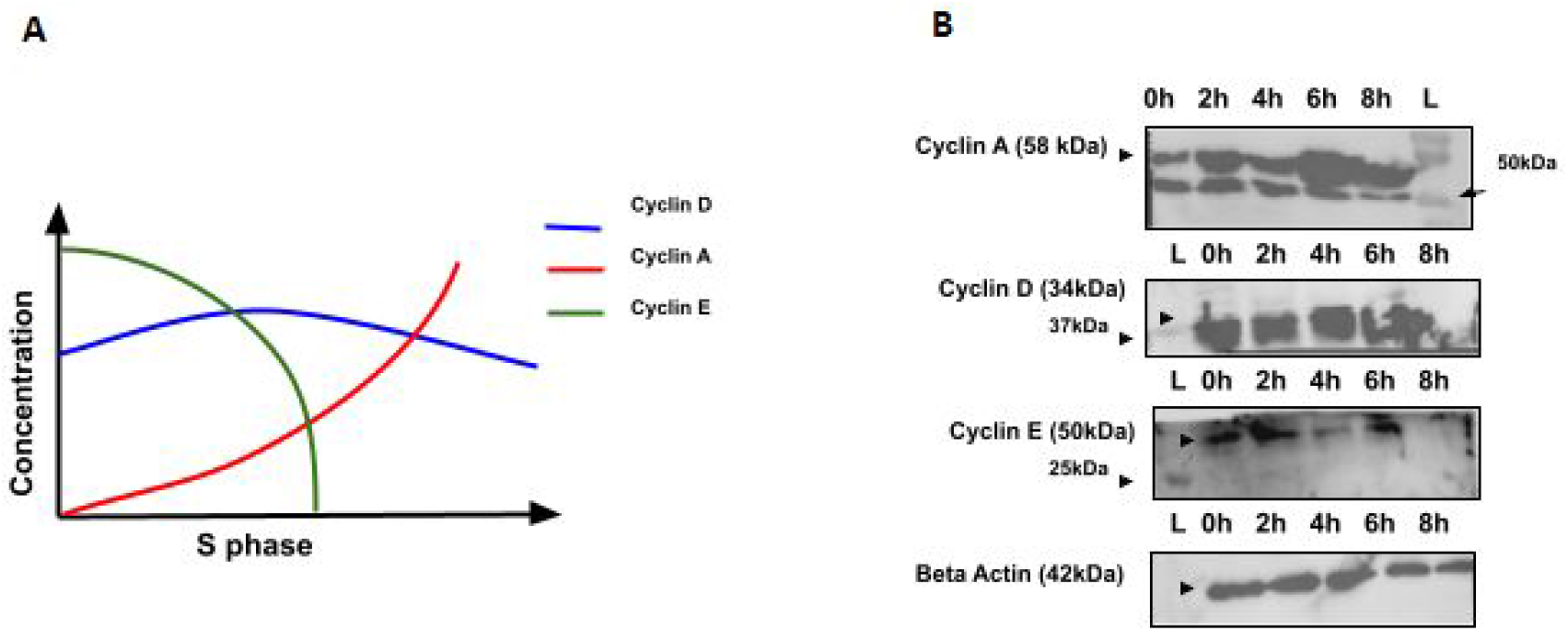
Assessment of cell synchronization by immunoblotting. (A) Representative figure of the cyclins in the S-phase. Cyclin A (red) increases in concentration as the cell progresses through the S-phase. Cyclin E (green) declines in the S-phase after activation of cyclin A. Cyclin D remains constant throughout the S-phase. (B) Western blot results of the amount of cyclins expressed in the HEK293T cells as a marker of cells progressing through S-phase after release of double thymidine block. Beta actin (42 kDa) is shown as a loading control.

**Fig 3:**
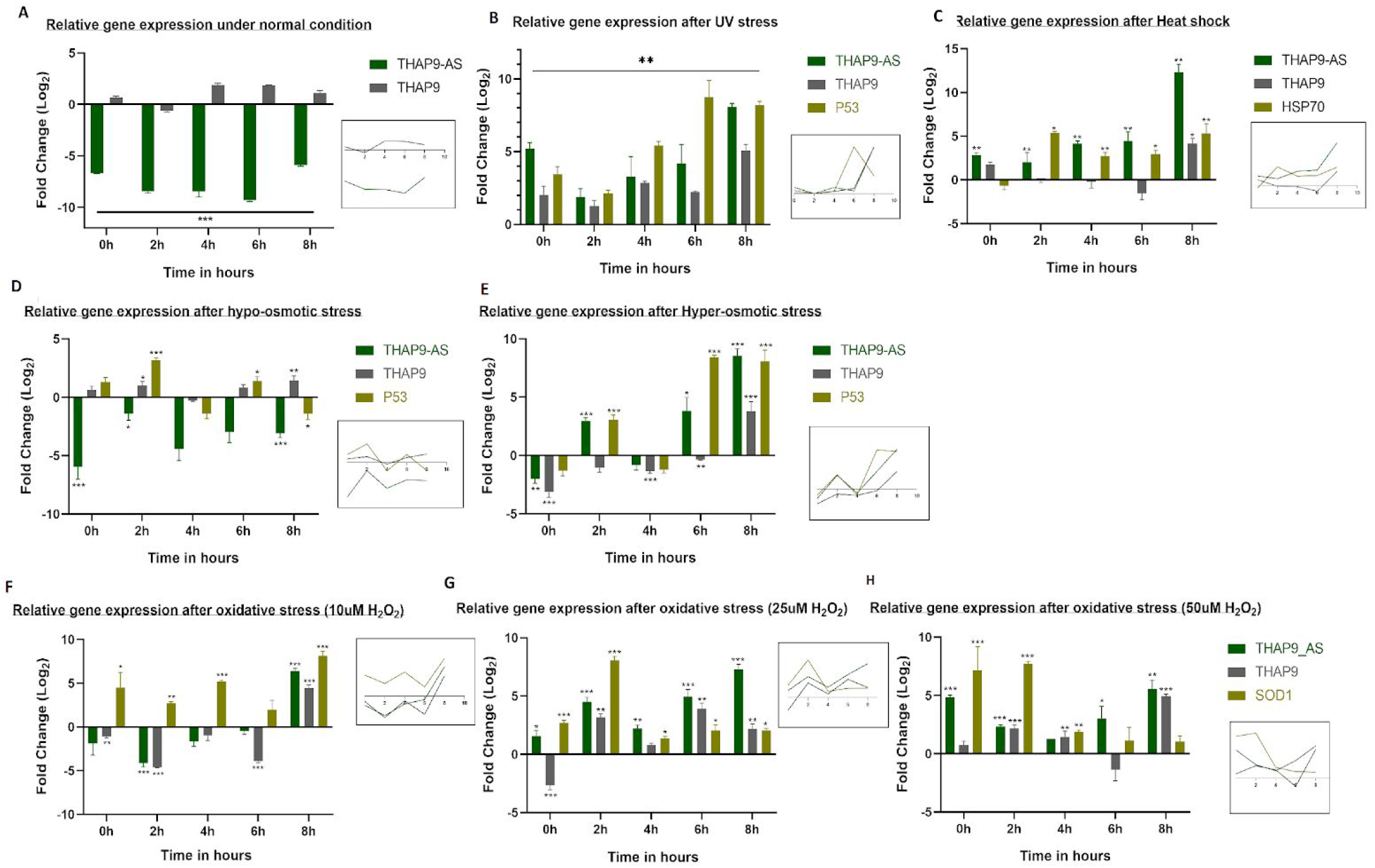
RT-qPCR analysis of THAP9 and THAP9-AS1 under stress conditions in the S-phase. Relative gene expression data represented as mean ± SEM (n=4, n=3 for oxidative stress) from 2 independent experiments. Data is normalized with HPRT1 gene expression data and represented as Log_2_ of fold change on y axis, time of sample collection on x axis. Upregulation of genes is represented on the positive y-axis and downregulation is shown on negative y-axis. Statistical significance was determined by two-tailed unpaired Student’s t test (*p<0.01, **p<0.001, ***p<0.0001).

### THAP9 and THAP9-AS1 exhibit opposite trends of gene expression after heat shock

To investigate whether heat has any effect on the gene expression of THAP9-AS1 and THAP9 we subjected the HEK293T cells to heat shock. We observed an overall downregulation of THAP9 mRNA expression throughout the S-phase and a sudden increase in expression in late S-phase (8 hour). Contrary to THAP9 expression, THAP9-AS1 transcript expression upregulates continuously from 2 hour to 8 hour in S-phase to a final 10 fold increase (p < 0.001) (Fig.3). HSP70 (Heat shock protein) which is a positive biomarker for heat stress also upregulates from mid to late S-phase (Fig 3).

### Relative gene expression of THAP9-AS1 changes in hypo-osmotic and hyper-osmotic states of cell

We observed that THAP9-AS1 exhibits differential gene expression under different types of osmotic stress. It is highly downregulated under hypo-osmotic conditions throughout the S-phase (p<0.001), with maximum downregulation in early S-phase (more than 5 fold decrease). On the other hand, THAP9’s gene expression does not change, except for a slight decrease in mid-S phase and an approximately two-fold increase in late S-phase (p<0.001). P53 was used as a positive marker for osmotic stress and maximum increase in its expression was observed in early S-phase (2 hour) (p<000.1). Conversely, under hyper-osmotic conditions, the expression of THAP9-AS1 increases in late-S phase (6-8 hour), after an initial downregulation in early S-phase (0h) (p<0.0001). THAP9-AS1 expression escalates to ~10 fold by the end of the S-phase. We noticed that its pattern of expression is similar to that of P53 (p<0.0001).

We did not observe much change in the overall THAP9 expression under hypo-osmotic stress with an exception during mid-S phase (4 hour), where THAP9 decreases. Under hyper-osmotic stress, THAP9 is highly downregulated in early and mid S phase but its expression surges in the end of S-phase (p<0.0001).

### Increase in concentration of H_2_O_2_ triggers an increase in the expression of THAP9-AS1

To induce oxidative stress, the cells were exposed to three different concentrations of hydrogen peroxide (H_2_O_2_), i.e., 10uM, 25uM and 50uM. At the lowest concentration of H_2_O_2_ (10uM), THAP9-AS1 and THAP9 are downregulated throughout the S-phase and increase to ~7 fold and ~5 fold respectively in the final stages of the S-phase (p<0.0001). When the concentration of H_2_O_2_ was raised to 25uM, both THAP9-AS1 and THAP9 is upregulated after 2hours in S phase. The expression of THAP9-AS1 stays higher than THAP9, although both the genes show a decrease in expression in the middle of the S-phase (4 hour). The highest transcript levels for THAP9-AS1 are recorded at 8 hour and for THAP9 at 6 hour.

At the highest concentration of H_2_O_2_ (50 uM), we observed that THAP9-AS1 exhibits an oscillatory pattern of expression. It decreases until the mid-S phase followed by an increase in expression until the end of the S-phase (p<0.001). THAP9, on the other hand, increases to ~2 fold in early-S phase and then downregulates at 6 hour followed by ~5 fold increase at 8 hour. Superoxide dismutase-1 (SOD1) gets activated in response to the oxidative stress and serves as a positive biomarker for the same. We observe that with increase in the concentration of H_2_O_2_, the upregulation of SOD1 gets triggered in early S-phase (0-2 hour) in comparison to the lower concentration of H_2_O_2_ (10uM) which leads to the highest expression of SOD1 in the end of the S-phase (8 hour).

## Discussion

Cells respond to external and internal stress via elaborate, multi-faceted systems and pathways. Most intrinsic stresses cause DNA lesions that are surveyed at DNA damage checkpoints during the cell cycle (42). Many transposons or ‘jumping genes’ are known to be activated under stress conditions which can lead to insertional mutagenesis and genomic instability (7, 43, 44).

Cancer cells exhibit an intrinsic stress due to altered metabolic activities (45). For example, a majority of tumor cells exhibit high oxidative stress to maintain high cell proliferation and facilitate the process of EMT (epithelial-mesenchymal transition) (46). Genome-wide studies of different types of cancers demonstrate that transposons are often responsible for oncogenic translocations (43, 44, 47). As a safety measure, transposons are often epigenetically silenced by repressive histone modifications or regulated post-transcriptionally by lncRNAs to counter ectopic expression of these dynamic sequences (7).

The activation of TEs under stress can be both beneficial or deleterious for the host (7). This is because the actual relationship between stress and TE expression is quite complex and varies depending upon the type of stress, type of organism and the epigenetic/genetic/metabolic changes drawn out by the stress (7). The regulation may be temporal or spatial wherein certain TEs can be turned on at specific stages of embryonic development or in specific cell types or tissues (7). Since stress induces both the defence response and the activation/deactivation of certain TEs within a host, it is hard to decipher the cause-effect relationship between the TEs and stress. Stress often results in double stranded DNA breaks which are detected and repaired at the DNA-integrity checkpoints of the cell cycle, by NHEJ (Non homologous end joining) when they occur in early S/G_1_ phase or Homologous repair when they occur in late S/G_2_ phase (21, 22). Interestingly, p53, which is known for its role in maintaining genomic stability during external stress and is mutated in almost all cases of cancer (23–25), may directly repress certain transposons (26).

LncRNAs which were long considered “junk” are now emerging to be important players in almost all aspects of cellular regulation (40, 48, 49) including cell proliferation in the cell cycle S-phase (39, 40, 49, 50) and regulation (e.g., by PANDA, lncRNA p21) of genes crucial to DDR (DNA damage response) (51–54). lncRNAs have been implicated in several diseases including cancer, diabetes and neurodegenerative disorders (55). Several lncRNAs have been identified as oncogenes or tumor suppressors which show cancer-specific upregulation (56). LncRNAs can interact with proteins and serve multiple roles like, decoys, scaffolds and guides (57). However, the role of several lncRNAs are poorly understood because their expression is highly specific to cell type, cellular stage, stimuli, experimental conditions (48, 58–61).

THAP9-AS1 is a recently annotated lncRNA and has been shown to be upregulated in pancreatic cancer and neutrophil apoptosis and downregulated in case of septic shock (35–37). In this study, we have quantified THAP9 and THAP9-AS1 transcript expression under various stress conditions in the S-phase of cell cycle. Under normal conditions, THAP9-AS1 is highly downregulated and THAP9 is upregulated in HEK293T cells (Fig. 3).

Ultraviolet rays are highly genotoxic and cause severe DNA damage leading to cell death. Genotoxic agents (like chemotherapy) are also used to treat cancer but therapy-resistant cancer cells are able to evade the damage by inhibiting tumor suppressor genes, increasing cellular growth factors and eluding cell cycle checkpoints (62). Several lncRNAs (NONHSAT1010169, GUARDIN, NEAT1) show changes in gene expression under genotoxic stress and are known to contribute to drug resistance in therapy-resistant cancer (50). P53 is responsible for genome defense in case of DNA damage and is known to be regulated by several lncRNAs (linc-p21, MEG3, TUG1, PANDA, PRAL and LED) in cancer and stress (50, 53, 56, 63, 64). We observed that genotoxic stress induces steady increase of THAP9-AS1 expression to a final ~10 fold upregulation by the end of S-phase (Fig 3A). Although its function is unclear but previous reports of its upregulation in pancreatic carcinoma suggests implication of THAP9-AS1 in carcinogenesis and DNA damage response.

LncRNAs also play an integral role in heat shock response. HSR1 (Heat shock RNA -1) plays an accessory role in forming the trimeric complex of HSF1, a heat shock protein for binding to DNA (65). Additionally, lncRNAs also regulate transcription of heat shock response genes (66). Here we report downregulation of THAP9 after a brief heat shock (Fig 3B), which might be a result of repression of general transcription under heat stress (66). Interestingly, on the other hand, THAP9-AS1 is upregulated by heat.

Osmotic stress affects the volume and tonicity of the cell and creates DNA lesions thereby eliciting a DNA damage response similar to that of UV stress (67). Hyper-osmotic stress is known to increase the expression of DNA damage responsive genes but not much is known about the participation of non-coding RNA in osmotic response in mammalian cells (68). In this study we have observed opposite effects on expression of THAP9-AS1 under osmotic stress. It shows downregulation under hypo-osmotic conditions but is upregulated in hyper-osmotic stress (Fig 3C, Fig 3D). A recent study suggests that hypo-osmotic stress cannot be linked to the misregulation of the cell cycle (69). This stands congruent to our findings where the expression of THAP9 and THAP9-AS1 under normal conditions seems similar to that recorded after hypo-osmotic stress.

Mitochondria, a key organelle involved in stress response, play a crucial role in maintaining cellular redox homeostasis by production of ROS (Reactive oxygen species). Oxidative stress, which occurs when ROS production surpasses the antioxidant defences, is known to damage DNA and disrupt mitochondrial function, as observed in cancer cells (70) as well as age related pathologies and cardiovascular diseases (71). Interestingly, THAP9 localizes in mitochondria, and its expression changes when subjected to oxidative stress using hydrogen peroxide. lncRNAs (MALAT1, NEAT1, H19, HULC) are upregulated in response to oxidative stress (50, 72). As we increase the concentration of hydrogen peroxide to trigger oxidative stress, the upregulation of THAP9-AS1 begins early in the S-phase.

In most reported sense-antisense gene pairs, the antisense gene can regulate the expression of the sense gene by RNA interference or by transcriptional interference, i.e, by competing for the enzymes essential for transcription on the shared promoter sites (33). *De novo* transcripts can arise in antisense configuration and are known to be positively correlated to their corresponding sense transcript under stress conditions (73). in most stress conditions the expression of both the transcripts increases. Given the overlapping promoters of THAP9 and THAP9-AS1, it can be hypothesized that the two genes may regulate each other’s expression or they share a common switch that contributes to the expression bias under certain stress conditions. Under basal conditions it is observed that THAP9 is preferentially transcribed whereas under stress conditions, THAP9-AS1 is transcriptionally favoured. This leads us to speculate that there exists a stress specific regulation of THAP9 and THAP9-AS1 gene-pair that switches to THAP9-AS1 under stress conditions. This also might explain the upregulation of THAP9-AS1 in cancer cells which are naturally stressed (35–37). Moreover, it is possible that both these genes play an unknown role in DNA repair and this might be responsible for their increased expression under certain stress conditions.

We also observe a periodicity in the expression of both THAP9 and THAP9-AS1 throughout the S-phase of the cell cycle, wherein, the amount of transcripts first decreases and then increases and vice versa (Fig 3B, 3D, 3F, 3G, 3H). The periodic gene expression of THAP9 and THAP9-AS1 resembles the periodicity pattern reported for the S-phase regulated expression cluster (74). The upregulated genes in S phase are also known to be involved in DNA damage response (74). The expression of THAP9-AS1 is much higher than THAP9 in all the stress conditions used in this study. The smaller length of THAP9-AS1 transcript in comparison to THAP9 facilitates faster expression and oscillations (75). One of the possible reasons for the same could be the less time required for lncRNA to respond to stress in comparison to a protein which requires additional steps of translation and folding.

## Conclusions

We show that THAP9-AS1 which is downregulated in normal state, turns out to be highly upregulated in different types of stress conditions. THAP9 on the other hand, exhibits a stress specific expression pattern. Furthermore, both THAP9 and THAP9-AS1 exhibit an oscillatory expression throughout the S-phase after subjecting the cells to stress. The pattern of expression of the two genes changes under various stresses, for eg, THAP9 is upregulated after exposure to UV rays and oxidative stress whereas it shows a downregulation under hyper-osmotic conditions and heat shock and a close to normal expression in hypo-osmotic state.

With emerging knowledge of sense-antisense gene pairs as a regulatory phenomenon in mammalian cells, it would be interesting to explore the relationship between THAP9 and THAP9-AS1. Although the function of both the genes remains elusive, this study has paved the way for exploring the function of THAP9 and THAP9-AS1 gene-pair in the context of DNA damage response.

## Acknowledgments

We would like to thank Dr. Rahul Kanadia, conversations with whom inspired this study and Dr. Umashankar Singh, Dr. Dhiraj Bhatia, Divyesh Patel, Manthan Patel, Subhamoy Dutta for their insights while conducting the experiments. Special mention to Pabba Maruthi Kumar for helping Meena in the preliminary experiments. We also acknowledge Dr. Umashankar Singh for generously sharing cyclin antibodies and the Himedia PCR machine (Insta Q96™ LA1012).

